# Genome-resolved metagenomics reveals metabolically versatile Actinobacteria with extensive biosynthetic capacity in Hawaiian steam vents

**DOI:** 10.64898/2026.09.14.751414

**Authors:** Shekhar Nagar, Chengxuan Zhang, Jimmy H. Saw

**Affiliations:** Department of Biological Sciences, The George Washington University, Washington DC, USA; Department of Zoology, Deshbandhu College, University of Delhi, New Delhi, India

**Keywords:** Actinobacteria, Hawaiʻi, ARGs, CO oxidation, fumaroles

## Abstract

Geothermally active lava caves and hydrothermal steam vents are chemically heterogeneous subsurface environments that harbor diverse microbial communities, yet the ecological and metabolic roles of Actinobacteria in these systems remain poorly characterized. Here, we reconstructed and analyzed 58 actinobacterial metagenome-assembled genomes (MAGs) representing five classes. Comparative genomic analyses revealed broad metabolic versatility, including pathways for amino acid biosynthesis and utilization, central carbon metabolism, fatty acid degradation, and potential chemolithotrophic processes involving carbon monoxide and sulfur compounds. The MAGs also encoded diverse carbohydrate-active enzymes and peptidases suggesting substantial capacity for complex organic carbon and protein utilization. Genome mining identified 219 biosynthetic gene clusters spanning terpenes, ribosomally synthesized and post-translationally modified peptides, nonribosomal peptide synthetases, polyketide synthases, and β-lactones, highlighting extensive secondary-metabolite biosynthetic potential. Antibiotic resistance-associated genes were also detected in several MAGs, with glycopeptide-resistance-associated *van* genes particularly prevalent among *Thermoleophilia*. Sequence similarity network analysis revealed that several *van*-associated proteins from *Thermoleophilia* and UBA4738 shared substantial sequence similarity with homologs from other environmental Actinobacteria, suggesting broad conservation of these protein families. Collectively, these findings reveal metabolically and functionally diverse Actinobacteria with the genomic potential to participate in carbon and nutrient cycling, microbial interactions, secondary metabolism, and antibiotic resistance in geothermally active ecosystems.

**Importance:** Actinobacteria are among the most extensively studied bacterial groups for their roles in terrestrial ecosystems and their capacity to produce bioactive compounds, yet their diversity and ecological functions in geothermal environments remain largely unexplored. Hydrothermal steam vents provide a unique setting in which steep chemical gradients may select for unusual combinations of metabolic, biosynthetic, and stress-response traits. Our genome-resolved analysis of Actinobacteria from Hawaiian geothermal habitats expands the known functional landscape of this phylum and reveals substantial diversity among lineages that are rarely represented in cultivated genome collections. The resulting genomes provide a framework for linking phylogenetic diversity with ecological function and for identifying candidate pathways underlying persistence in chemically dynamic environments. More broadly, this study demonstrates the value of genome-resolved approaches for uncovering the functional potential of microbial lineages that remain inaccessible to conventional cultivation and provides targets for future experimental investigation.

## Introduction

In volcanic ecosystems, microbes often exhibit strong interdependence, especially in their interactions with chemolithotrophs that use reduced compounds such as nitrate and sulfate components as electron acceptors in basalt as energy sources (1, 2). Fumaroles and geothermally active lava caves are dynamic and extreme environments that form when steam and volcanic gases escape through cracks in basalt deposits, creating vents (3, 4). Unexpectedly, high microbial diversity has been observed in oligotrophic volcanic features in Iceland, the Azores, the Canary Islands, and the Hawaiian (4, 5). Previous research over these sites has determined that subsurface microbial consortia develop localized, site-specific microbial–mineral interactions over time. This occurs as bacteria acclimate and adapt to specific mineral chemistry, nutrient availability, and atmospheric conditions on a microscopic scale (6–8). Large-scale ecological factors, such as the age and extent of basalt weathering, as well as stochastic and deterministic processes, may also affect bacterial diversity in lava caves, shaping microbial communities over time. Experimental studies on biofilm-mineral interactions in caves or basalts have been conducted, but our understanding of these interactions is still limited (9). These studies have raised many unanswered questions about how microbial diversity and ecology may differ in hydrothermal systems that appear due to the Hawaiʻi volcanic hotspot.

Actinobacteria are a widespread phylum with higher guanosine-cytosine (65-75% G + C) contents that thrive in nearly every environment, ranging from soil and marine habitats to more unexpected settings such as fossils, caves, flora and rhizosphere (10–12). Since the discovery of antibiotics in the 1940s, members of Actinobacteria have garnered significant attention, especially those that belong to the Actinomycetes class, which are recognized as key sources of therapeutic pharmaceuticals (10). Cave Actinobacteria are also intriguing due to the fact that caves are essentially oligotrophic habitats resembling extreme conditions with low nutrient availability and primary productivity (13, 14). These conditions probably constrain microbes to utilize distinct metabolic pathways, such as those involved in biomineralization, rock weathering, secondary metabolite production, and antibiotic resistance (15, 16). These Actinobacteria are also important sources of novel primary and secondary metabolites that can be harnessed in biotechnology or pharmaceutical industries (10). Over the past two decades, non-streptomycete actinomycetes (rare actinomycetes) have made a significant impact, accounting for a large percentage of all known antibiotics (17). A deeper understanding of the diversity and distribution of Actinobacteria can offer valuable insights into microbial ecology and aid in the discovery of novel bioactive compounds, including the role and diversification of antibiotics targeted sites found in less studied classes such as *Actinomycetia*, *Thermoleophilia*, and *UBA4738* (18, 19).

While the Actinobacteria have been studied extensively since the days of Selman Waksman, much less is known about the lineages that are found in extreme habitats, such as those found in volcanic fumaroles of Hawaiʻi. In addition, their metabolic potentials for other biological processes are also not well known. In this study, we employed genome-resolved metagenomics to explore the role of Actinobacteria in fumarole-associated biofilms and soil that are found within various volcanic regions of the Big Island (also simply known as Hawaiʻi). We constructed metagenome-assembled genomes (MAGs) from these samples and investigated the metabolic potential and diversity of secondary metabolites predicted from the MAGs.

We recovered 58 high- and medium-quality MAGs representing five classes (Acidimicrobiia, Actinomycetia, CALGFH01, Thermoleophilia, and UBA4738) and characterized their phylogenetic relationships and functional repertoires. The recovered genomes revealed extensive metabolic flexibility, combining pathways for heterotrophic utilization of amino acids, carbohydrates, lipids, and other organic substrates with genomic potential for chemolithotrophic processes involving carbon monoxide and sulfur compounds. Their diverse carbon-, nitrogen-, sulfur-, and iron-cycling capabilities further suggest that these lineages may contribute to elemental transformations across the steep physicochemical gradients characteristic of geothermal habitats. The genomes also contained extensive repertoires of carbohydrate-active enzymes and peptidases, together with diverse biosynthetic gene clusters encoding terpenes, RiPPs, NRPSs, PKSs, and β-lactones, indicating substantial potential for complex carbon utilization and secondary metabolism. Notably, antibiotic resistance determinants were widespread, with glycopeptide-resistance-associated *van* genes particularly enriched among Thermoleophilia, and sequence similarity analyses revealed conservation of several resistance-associated protein families across environmental lineages. Collectively, these findings reveal previously undercharacterized Actinobacteria with broad metabolic, biosynthetic, and resistance-associated potential and provide new insights into their possible ecological roles in geothermally active terrestrial ecosystems.

## Materials and Methods

### Sampling sites and processing

Site description, sample collection, and sequencing have been described in detail in a previous study (20). Briefly, samples were collected from steam vent associated geothermal features within the Hawaiʻi Volcanoes National Park or the East Rift Zone, DNA extracted in the lab at George Washington University, and sequenced at Oregon State University (20). The samples came from three different geothermal features: steam pits, steam walls, and cave soil. Each of these samples was assigned an identification code consisting of two parts - the former is a letter code describing the site of collection and the latter is a number unique to the sample (Table 1).

**Table 1.** Description of the sample features along with *in situ* temperatures measured at sites.

| Site | Sample | Temp (°C) | Description |
| --- | --- | --- | --- |
| Kilauea caldera | HW02 | 25 | Cave soil in Kilauea caldera |
| Kilauea caldera | HW12 | 25 | Cave soil in Kilauea caldera |
| Steam pit | P23_25 | 38 | harden purple biofilm inside a small cave |
| Steam wall | PDB5 | 43 | dark purple/grey dry biofilms near shaded regions of a group of steam vents |
| Steam wall | S13_15 | 43 | dark purple/grey dry biofilms near shaded regions of a group of steam vents |
| Steam wall | S18_20 | 55 | snottites under ceiling inside a small steam cave |
| Steam wall | S22_25 | 47-51 | hanging biofilms along rock walls near a group of steam vents |
| Steam wall | S29_31 | 47 | hard microbial mats with plant-like features near a group of steam vents |
| Steam wall | S2_4 | 43 | hanging biofilm on mountain-side of a group of steam vents |
| Steam wall | S34_36 | 43 | tofu-like microbial mats near shaded regions of a group of steam vents |
| Steam wall | S37 | 69 | green biofilm inside a hot steam vent on the ground |
| Steam wall | S7_8 | 50 | cream to green biofilms on exposed walls of a group of steam vents |
| Kilauea caldera | W1_4 | 34 | dry and crusty biofilms on a cave wall in Kilauea caldera |

### Metagenome assembly and binning

Detailed metagenome assembly and binning procedures were described in a previously published study (20). Briefly, BBDuk was used to trim raw metagenomic sequences to remove adapter and poor-quality sequences and assembled using metaSPAdes v3.15.4 (using k-mers of 21, 33, 55, and 77) (21). Trimmed reads were mapped back to resulting contigs using BBWrap v38.87 (sourceforge.net/projects/bbmap/) (22) and underwent binning using Metabat2 v2.15 (23). Samples that originated from the same features were combined and co-assembled (for example, S34, S35, and S36 were co-assembled and this co-assembly is named S34_36). CheckM was used to assess genome completeness and contamination and MAGs adhering to the MIMAG criteria (24), specifically those with ≥ 50% completeness and ≤ 5% contamination, were further analyzed.

### Identification, Classification and Phylogenetic Analysis

Based on MIMAG criteria (25), reconstructed MAGs were assigned as either HQ (High-Quality) or MQ (Medium-Quality). The ANI similarity amongst genomes present in GTDB database (release 214) and the MAGs was evaluated by FastANI (26) integrated in GTDB-tk (27). Taxonomic classification and identification of the MAGs was done using the Genome Taxonomy Database Toolkit (GTDB-tk) v2.1.1 with database release 214 (28, 29). Further, phylogenetic reconstruction (Bootsrap=100) was performed by using DIAMOND (30) for mapping, MUSCLE (31) for Multiple Sequence Alignment, and FastTree v2 (32) for ML-based approach tree construction. The MAGs were first annotated using Prokka (33) along with the UniProt database (34) and mapped based on marker genes. Finally, the Maximum-likelihood approach was used to infer the phylogenetic tree using PhyloPhlAn v3.0 (35). Resulting tree was visualized and exported using the ITOL v6.9.1 (36).

### Metabolic pathways reconstruction

Gene prediction of the MAGs were done with prodigal (37) followed by metabolic pathway analysis using Metabolic and Biogeochemistry Analyses in Microbes (METABOLIC) v4.0 pipeline (38). The algorithm used in METABOLIC-G.pl script includes utilization of HMM (Hidden Markov Models)-based metabolic databases that include KOfam (39, 40), Pfam (41), and TIGRfam (42). To estimate the homology of query proteins with a highly curated set of protein motifs in databases, -m-cutoff was set to 0.75. The resulting categorized pathways and gene ontologies of functional traits present in the MAGs were visualized in R Statistical Software (v4.1.2) (43). The proteins identified from the searches were also recruited against dbCANfam hmm database (CAZymes gene clusters) to define their carbohydrate active enzymatic signatures using dbCAN2 (44). The structural, functional and hierarchical based classification of annotated proteins was performed by clustering the proteins into peptidase clans/families using MEROPS HMM database (45).

### Nutrient cycling and Biosynthetic gene clusters

To estimate the biogeochemical nutrients the annotated proteins were searched against an open-source pipeline Multigenomic Entropy Based Score (MEBS) (46) that has been validated with a dataset comprising 2,107 unique microbial genomes from RefSeq and 935 metagenomes from MG-RAST. It is designed to evaluate, compare, and quantify biogeochemical cycles and calculate the relative entropy-based scores to assess the abundance of key ecological elements—Nitrogen (N), Carbon (C), Sulfur (S), Iron (Fe) and Oxygen (O) within an ecosystem (46). The hmmsearch tool integrated in MEBS employs a stringent false discovery rate (FDR) threshold of 0.001 against the protein domains of Pfam v3.0 database associated with N, C, S, Fe and O cycles. The coding sequences of each MAG were also examined using the antiSMASH v7.0 software (47) to search for biosynthetic gene clusters (BGCs) through HMMs. The antiSMASH software effectively identifies gene clusters that encode secondary metabolites by leveraging its comprehensive database of annotated domains and modules across a wide range of chemical classes.

### Antibiotic resistance genes

To determine the antimicrobial resistance across the recovered genomes we aligned the proteins of MAGs with the predicted Comprehensive Antibiotic Resistance Database (CARD v3.2.4, https://card.mcmaster.ca July, 2022) (48) using Resistance Gene Identifier (RGI) and ResFinder tools (49, 50). Then, the pipeline ARGs-OAP v3.0 (51) was used with two stage procedure for identification and quantification of ARGs with the help of proteins that were aligned with the structured ARG database (SARG) v3.0 (51) by BLAST+ v2.12.0 (similarity ≥ 80%, *e* value ≤ 1e − 7 and query coverage ≥ 75%). The SARG database is a curated database that consists of 713 sequences from CARD which includes beta lactam (230), aminoglycoside (157), quinolone (106), MDR (99), macrolide-lincosamide-streptogramin (MLS) (96), vancomycin (73) and chloramphenicol (35) and other resistance types. The identified ARGs in MAGs were visualized as circular ideograms and displayed in Circos table viewer v0.63-10 (52).

### Sequence similarity Analyses

ARG proteins identified in *Thermoleophila* and *UBA4738* classes by RGI and ARGs-OAP pipelines were merged and built into searchable database using *makedb*. They were searched against proteomes of related actinobacterial classes found in GTDB by BLASTp v2.15.0. The ARGs that showed >50% in identity and more than 90 amino acid residues in alignment lengths were taken into consideration to estimate the enzymatic sequence similarity networks (SSNs). Proteins found in MAGs belonging to *Thermoleophila* and *UBA4738* were further compared with reference genomes from GTDB to investigate the diversification of all abundant ARGs.

All resulting proteins found from the searches were filtered to remove identical sequences (100%) using CD-HIT (53) and The Enzyme Function Initiative-Enzyme Similarity Tool (EFI-EST) was used to calculate independent pairwise relationship of the enzyme families (54). Sequence similarity networks (SSNs) were built with a threshold alignment score of 50% and visualized using Cytoscape v3.7.2 (55). For the ARGs sequence networks, a cut-off value of 1e^-30^ was applied, based on an analysis of alignment length trends at various e-values. The average number of neighbors or degree for a node or sequence was calculated as k = 2K/N where K denotes the total number of edges and N denotes the total nodes. To estimate the diversity/similarity among sequences, the density of networks i.e. the fraction of all edges in the similarity networks was also calculated as D= 2K/N(N-1). These proteins were also partitioned in accordance with their source of isolation, summarized through NCBI datasets command-line tool (56).

## Results

### Phylogeny of novel genomes and distribution of Actinobacteria

Based on the MIMAG criteria (24), we analyzed 58 MAGs that fall into medium-quality or high-quality categories. These 58 Actinobacterial MAGs were classified into 5 classes: *Acidimicrobiia* (n=19), *Actinomycetia* (n=4), *CALGFH01* (n=7), *Thermoleophilia* (n=24), and *UBA4738* (n=4) (Figure 1). We were able to classify 39 of the MAGs up to the genus level (Supplemental Table S1). Here, most of MAGs (n=35) were high-quality genomes (completeness ≥ 90%; contamination ≤ 5%) with three 100% complete genomes from the genus *PALSA-610* (c_*Acidimicrobiia*) while other 23 were medium-quality genomes (completeness ≥ 50%; contamination ≤ 5%). Maximum-likelihood phylogenetic analysis based on a conserved set of marker genes reconstructed the evolutionary relationships among the 58 actinobacterial MAGs (Figure 1). The phylogeny recovered five distinct actinobacterial classes: *Acidimicrobiia*, *Actinomycetia*, *CALGFH01*, *Thermoleophilia*, and *UBA4738*. Members of *Acidimicrobiia*, *Actinomycetia*, *CALGFH01*, and *UBA4738* each formed well-defined monophyletic clades. In contrast, *Thermoleophilia* was resolved into two major lineages. One lineage comprised 12 MAGs assigned to the family *Gaiellaceae*, whereas the second contained 11 MAGs belonging to the order *Solirubrobacterales*, including members of the families *Solirubrobacteraceae*, *Thermoleophilaceae*, and family 70-9. The genome W1_4_bin_71, assigned to the order *Miltoncostaeales*, also clustered within the *Thermoleophilia* lineage.

**Figure 1.**
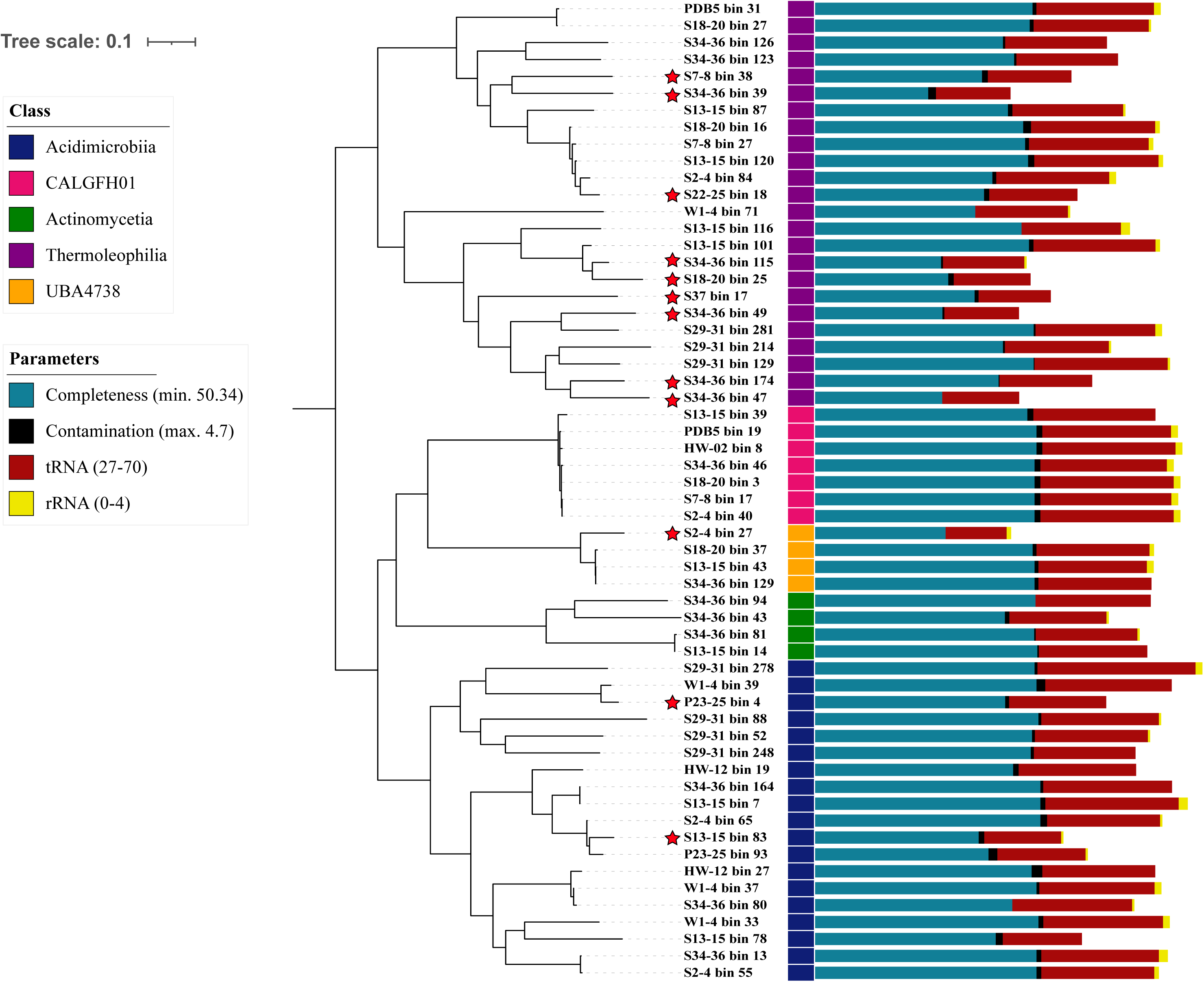
Phylogenomic tree of 58 actinobacterial MAGs. Color strips next to MAG ids show five classes: *Acidimicrobiia* (n=19), *Actinomycetia* (n=4), *CALGFH01* (n=7), *Thermoleophilia* (n=24), and *UBA4738* (n=4). Stacked histograms show features such as completeness (50.34-100 %; blue), contamination (0 - 4.7 %; black), number of tRNA copies (red), and number of rRNA copies (yellow). The genomes marked with red star highlights the MAGs with less than 70% ANI values to closest reference genomes in the GTDB database.

### Metabolic reconstruction (KEGG)

Metabolic reconstruction of the 58 actinobacterial MAGs revealed broad biosynthetic and catabolic capabilities (Figure 2). The KEGG analysis revealed a large number of genes involved in various pathways as follows: amino acid synthesis and metabolism (n=868), central carbohydrate metabolism (n=640), cofactor and vitamin metabolism (n=523), lipid metabolism and degradation (n=115), ATP synthesis (n=131), aromatic degradation and drug resistance (n=115), carbon fixation, nitrogen, sulfur metabolism (n=97). Nearly all MAGs encoded complete pathways for phosphoribosyl pyrophosphate (PRPP) biosynthesis (M00005), pentose phosphate metabolism leading to ribose-5-phosphate (M00007), arginine biosynthesis (M00844), and branched-chain amino acid biosynthesis, including valine and isoleucine (M00019 and M00570). Approximately 75% of the members have complete metabolic pathways that are involved in glycolysis, adenine biosynthesis, cytochrome c oxidase, pyridoxal phosphate biosynthesis, lysine biosynthesis via succinyl-DAP, and threonine biosynthesis (Figure 2A).

**Figure 2.**
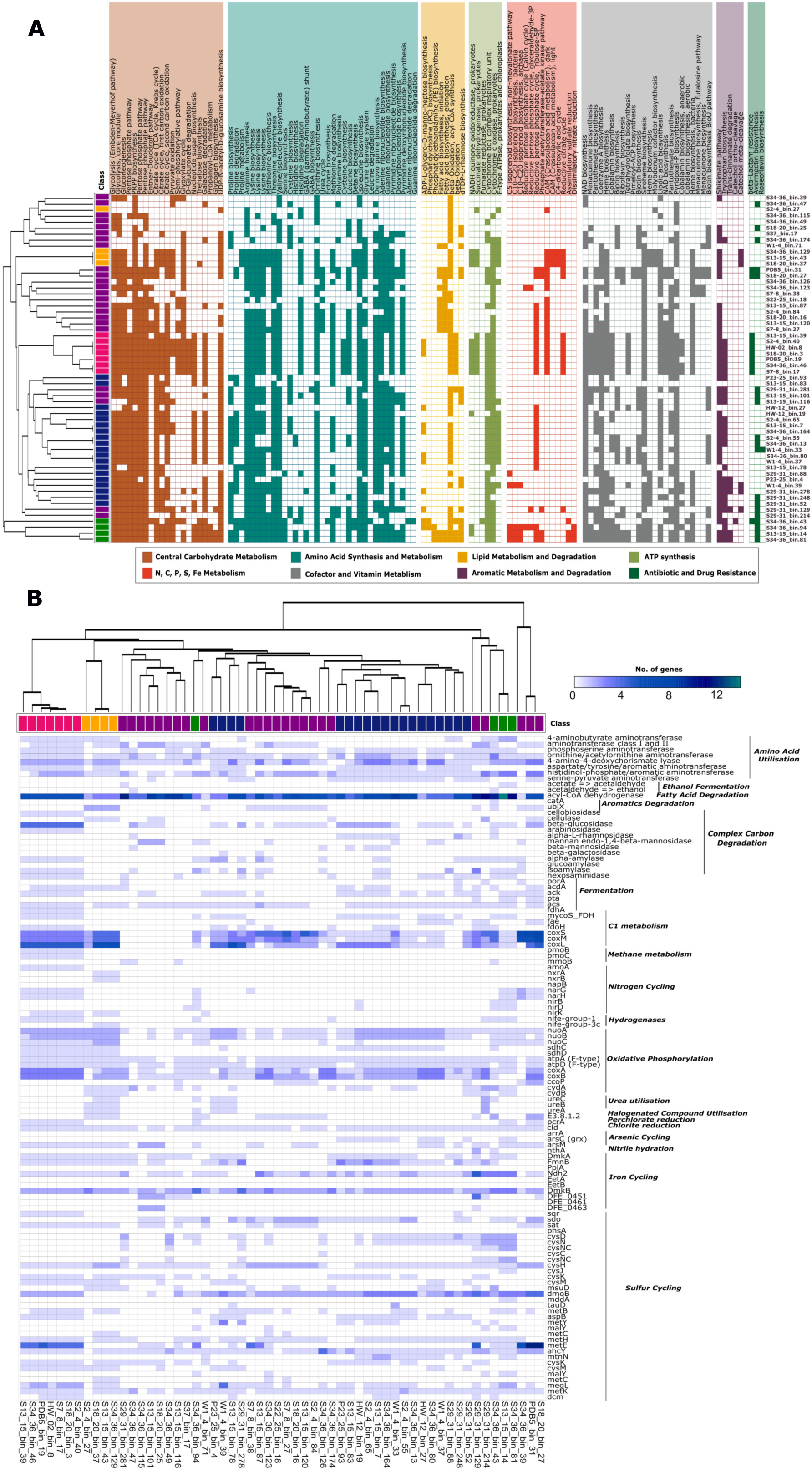
Basic metabolic profiling based on number of KEGG orthologs found. (A) Presence/absence of essential pathways involved in the metabolism of amino acids, lipids, central carbon and others. (B) The heatmap shows the copy number of gene clusters found in all metabolic pathways present in the actinobacterial MAG.

Several lineage-specific metabolic features were also observed. Complete reductive tricarboxylic acid (rTCA) cycle modules (M00173) were detected exclusively in members of the class CALGFH01. W1-4_bin.33 that belongs to Acidimicrobiia class might be able to synthesize roseoflavin, an antibiotic, from Flavin Mononucleotide (M00890). In addition, S34-36_bin.43 that belongs to Actinomycetia class encoded genes for KEGG modules (M00033; M00036; M00546; M00150; M00568) involved in ectoine biosynthesis, leucine degradation to acetyl-CoA, purine degradation via urea, fumarate reduction, and catechol degradation, indicating expanded metabolic versatility. Genes associated with fatty acid degradation (acyl-CoA dehydrogenase), sulfur metabolism (*dmoB*, *metE*, *cysH*, *ahcY*), oxidative phosphorylation (*coxA*, *coxB*), C1 metabolism (*coxS*, *coxM*, *coxL*), and amino acid utilization (branched-chain amino acid transferase) were also found in various abundances within the MAGs.

### Carbohydrate-active enzymes (CAZymes)

Annotation against the dbCAN database identified 1,432 carbohydrate-active enzymes (CAZymes) belonging to 67 CAZy families across the 58 MAGs. Members of the class Thermoleophilia encoded the largest CAZyme repertoires overall, whereas Acidimicrobiia contained the highest abundance of α-amylases (GH13). The most abundant CAZy family was GH13 (α-amylases), with 191 copies detected across the dataset, indicating a widespread capacity for starch and glycogen degradation. GH13 enzymes were identified in all five actinobacterial classes and were particularly enriched in Acidimicrobiia. Members of Thermoleophilia were instead enriched in GH39 α-L-iduronidases, enzymes involved in glycosaminoglycan degradation, suggesting an enhanced capacity to utilize complex host- or biofilm-derived polysaccharides. At the genome level, the UBA4738 genome S29-31_bin.129 encoded the largest CAZyme repertoire (49 genes), including numerous GH13 α-amylases and GH39 α-L-iduronidases. The Thermoleophilia genome S29-31_bin.214 contained eight GH79 β-glucuronidases, further supporting the ability of this lineage to degrade complex carbohydrates.

### Peptidases (MEROPS)

Screening against the MEROPS database identified 2,417 peptidases representing 75 peptidase families. Acidimicrobiia and Thermoleophilia encoded the largest numbers of peptidases, followed by Actinomycetia, CALGFH01, and UBA4738. The most abundant peptidase family was S33 (prolyl aminopeptidases), with 377 copies detected across the dataset (Figure 3A). These enzymes remove N-terminal proline residues from peptides and likely contribute to protein turnover. The highest copy numbers were observed in the Actinomycetia genomes S13_15_bin.14 and S34_36_bin.81. Members of Thermoleophilia were particularly enriched in the C26 family of γ-glutamyl hydrolases, enzymes that hydrolyze γ-linked glutamate residues from polyglutamylated substrates such as folates (57). This enrichment suggests an enhanced capacity for folate metabolism and amino acid recycling within this lineage.

**Figure 3.**
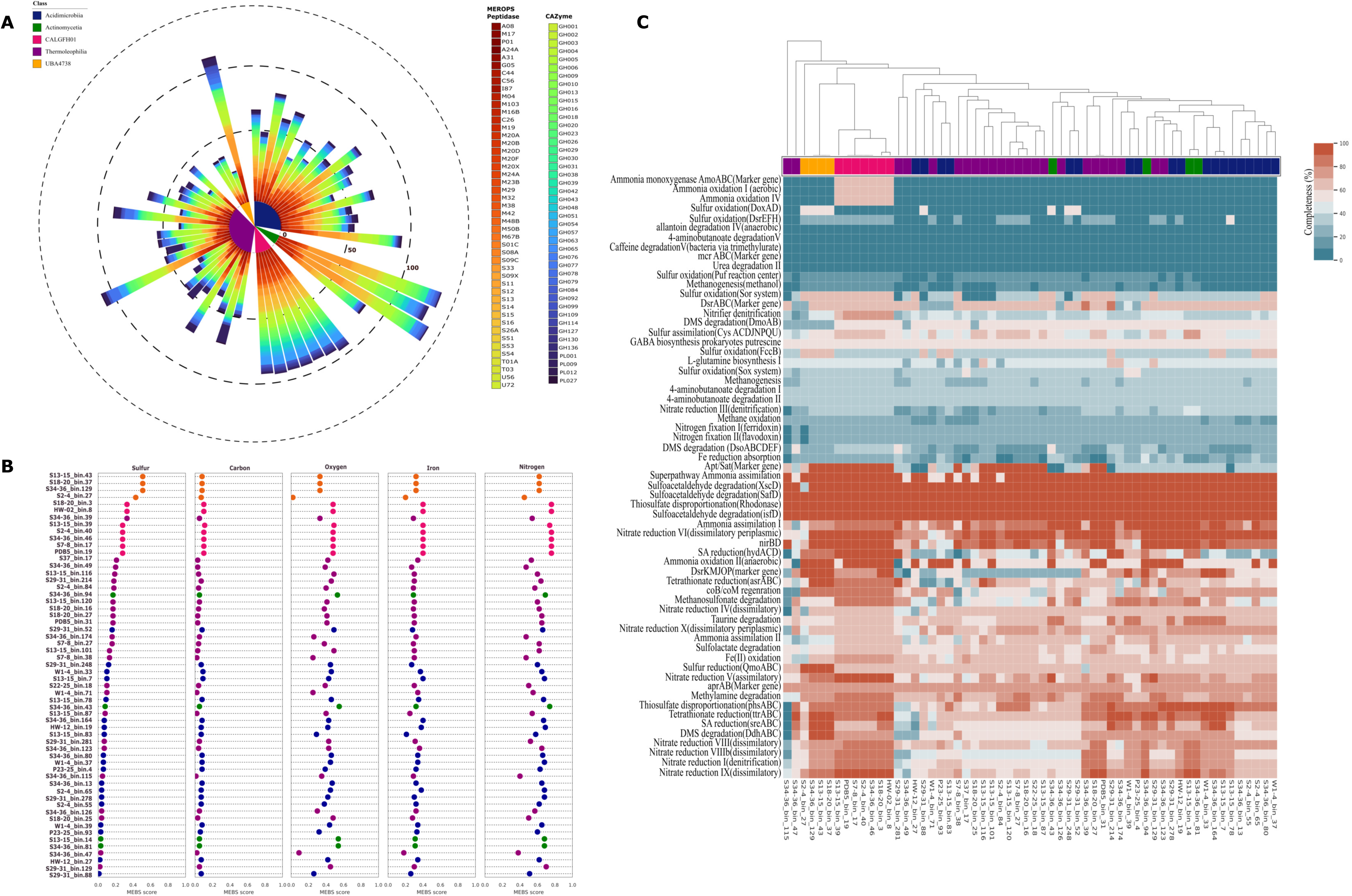
Carbohydrate and biogeochemical cycling potential of the 58 Actinobacteria MAGs. (A) dbCANfam showed the presence of 67 families of CAZy (Carbohydrate-Active EnZymes database). (B) Entropy scores (H’) of N, S, C, O and Fe estimated across all MAGs and MAGs sorted from lower to higher sulfur entropies to assess relative biogeochemical cycling potential of them. (C) Metabolic completeness of essential biogeochemical pathways shown in color gradients – more completed pathways are shown in red color gradient and less completed pathways are shown in blue color gradient. The color strips on top of the heatmap correspond to the phylum-level classification of MAGs as shown in Figure 1.

### Biogeochemical cycling potential

We assessed the potential of the 58 actinobacterial MAGs to participate in biogeochemical cycling of carbon, nitrogen, sulfur, oxygen, and iron using the MEBS framework. Overall, the MAGs encoded diverse sets of genes associated with carbon, nitrogen, sulfur, and iron transformations, although the distribution of these functions varied among taxonomic classes (Figure 3B, C). Members of CALGFH01 exhibited relatively high entropy scores for carbon, oxygen, iron, and nitrogen cycling, with mean scores of 0.11, 0.48, 0.40, and 0.75, respectively. In contrast, UBA4738 showed the highest mean sulfur-cycling entropy score (0.49), whereas sulfur-cycling scores were lower in Thermoleophilia (0.13), Acidimicrobiia (0.06), and Actinomycetia (0.07). Genes associated with several core nutrient-cycling pathways were broadly distributed across the MAGs. Pathways for ammonia assimilation, thiosulfate disproportionation, and sulfoacetaldehyde degradation were detected across all five classes. In contrast, genes associated with ammonia oxidation, including the AmoABC complex and ammonia oxidation modules I, II, and IV, were detected primarily in CALGFH01. Sulfur oxidation and sulfur-reduction pathways, including the Sor system, Apt/Sat, DsrEFHABCKMJOP, HydACD, and SreABC, were particularly prevalent among UBA4738 and CALGFH01 MAGs.

Nitrogen-cycling genes were also widespread. Nitrogen fixation pathways were more complete in CALGFH01 (63.6–72.7% of pathway completeness) compared to other classes (36.4– 45.5%). The *nirBD* gene cluster was detected across all five classes, with pathway completeness ranging from 40–100% in Acidimicrobiia and Thermoleophilia, 60–100% in UBA4738, and up to 100% in Actinomycetia and CALGFH01. Most MAGs also encoded the DdhABC pathway associated with dimethyl sulfide degradation and methanethiol production. The AC-56 genome S13-15_bin.101 was an exception, lacking detectable nitrate reduction, nitrogen fixation, and DdhABC pathway components. Iron-cycling potential also differed among lineages. Genes associated with dissimilatory iron reduction were more prevalent in Thermoleophilia, whereas genes associated with Fe (II) oxidation were particularly abundant in CALGFH01, with pathway completeness reaching 77.8%. Together, these results indicate that the actinobacterial MAGs possess diverse metabolic capacities that could contribute to carbon, nitrogen, sulfur, and iron transformations in hydrothermal steam-vent environments.

### Secondary metabolite biosynthetic potential

Genome mining identified extensive secondary metabolite biosynthetic potential across the 58 actinobacterial MAGs. A total of 219 biosynthetic gene clusters (BGCs) were detected, including clusters associated with terpenes, ribosomally synthesized and post-translationally modified peptides (RiPPs), nonribosomal peptide synthetases (NRPSs), polyketide synthases (PKSs), and β-lactones (Figure 4A). Terpene-associated BGCs were the most abundant (n = 70), followed by RiPPs (n = 57), NRPSs (n = 32), PKSs (n = 20), and β-lactones (n = 17). The distribution of BGCs differed among taxonomic classes. Actinomycetia accounted for approximately 75% of the NRPS-associated BGCs, with particularly high representation among the Mycobacterium-affiliated MAGs, whereas CALGFH01 contained approximately 65% of the PKS-associated BGCs. RiPP-associated BGCs were detected primarily in Thermoleophilia (n = 24), Acidimicrobiia (n = 25), and Actinomycetia (n = 8), but were rare or absent in CALGFH01 and UBA4738. Several less abundant BGC classes were also detected. These included type III polyketide synthases and HglE-KS-associated clusters in CALGFH01; RRE-containing, type II polyketide synthase, and LAP-associated clusters in Thermoleophilia; arylpolyene and ranthipeptide clusters in Acidimicrobiia; and indole- and oligosaccharide-associated clusters in UBA4738. Collectively, the broad distribution of BGCs across the five actinobacterial classes indicates substantial biosynthetic potential and suggests that hydrothermal steam-vent Actinobacteria may contribute to the chemical diversity of these microbial communities.

**Figure 4.**
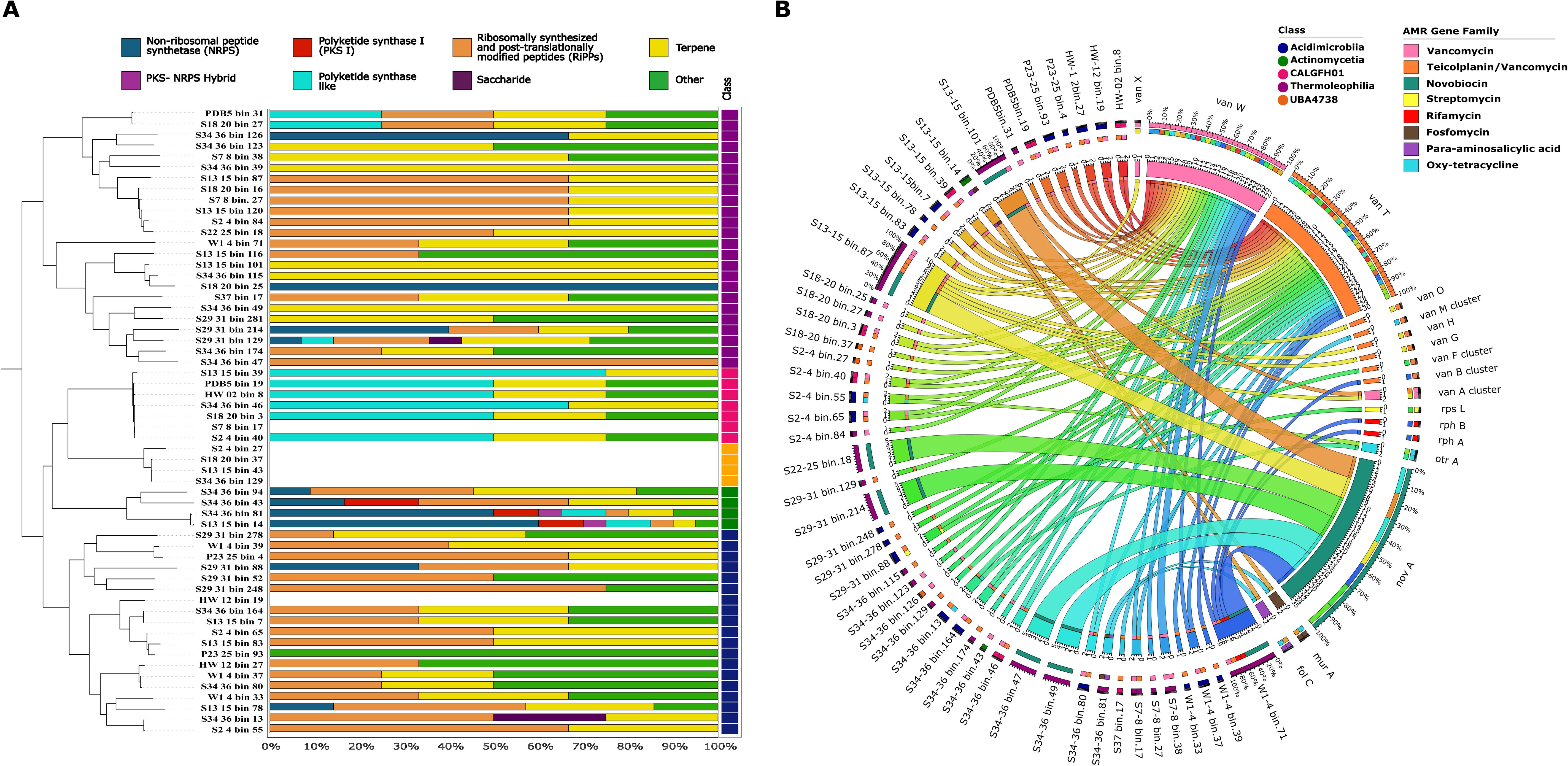
Secondary metabolites and antibiotic gene repertoire in the actinobacterial MAGs. (A) Percentage distribution of biosynthetic gene clusters found in the 58 MAGs. (B) Circos layout visualizing the abundance of 119 ARGs in 50 MAGs that belong to different antimicrobial resistance gene families. The color strips in the outermost circle represent the classes of MAGs (5-12 o’clock positions) and AMR gene families (12-5 o’clock positions). Cords connecting MAGs to AMR families are drawn if a certain AMR family is detected in each MAG and percentages of their distributions are shown where needed.

### Antibiotic resistance gene repertoire

We characterized the antibiotic resistance gene (ARG) repertoire of the 58 MAGs using ARGs-OAP and RGI. ARGs were detected in 50 MAGs (86%), representing eight resistance classes and 17 ARG subtypes, with a total of 116 predicted resistance determinants (Figure 4). The most frequently detected genes were *novA* (n = 35), *vanT* (n = 32), and *vanW* (n = 31). ARG abundance varied among taxonomic classes, with Thermoleophilia exhibiting the highest overall ARG representation. The Thermoleophilia MAGs belonging to g PALSA-612 (S13-15_bin.87 and W1-4_bin.71) contained the largest ARG repertoires. Thermoleophilia contained 61 ARG subtypes, compared with 31 in Acidimicrobiia, 12 in CALGFH01, eight in Actinomycetia, and four in UBA4738. When normalized to estimated 16S rRNA gene copy number, Thermoleophilia also exhibited a relatively high ARG density. Glycopeptide-resistance determinants were particularly widespread. *vanT* was detected in 30 MAGs spanning Acidimicrobiia, Actinomycetia, Thermoleophilia, and UBA4738, whereas *vanW* was detected in 26 MAGs belonging to Acidimicrobiia, Actinomycetia, and Thermoleophilia.

Within Thermoleophilia, glycopeptide-resistance determinants were dominated by *vanABFGHMOTX* and *vanW*. The MAG W1-4_bin.71 additionally contained *rphA* and *rphB*, which were identified as rifamycin-resistance determinants. The *novA* gene was detected exclusively among Thermoleophilia MAGs in this dataset, occurring in seven MAGs and representing a major component of their predicted resistance repertoire. Other resistance determinants were detected in individual lineages. The Acidimicrobiia MAG S29-31_bin.88 contained *rpsL*, a determinant associated with streptomycin resistance, whereas two JACDBD01-affiliated MAGs contained *otrA*, associated with oxytetracycline resistance. Within Actinomycetia, *folC* and *murA* were detected in two *Mycobacterium*-affiliated MAGs and were associated with resistance to para-aminosalicylic acid and fosfomycin, respectively.

### Sequence similarity networks reveal conserved glycopeptide-resistance determinants

To investigate the evolutionary conservation and diversification of glycopeptide-resistance determinants identified in Thermoleophilia and UBA4738, we compared the predicted ARG proteins with reference sequences using sequence similarity networks (SSNs) (Figure 5). Among the Thermoleophilia MAGs, *rphA/rphB* sequences showed limited similarity to reference sequences from other Thermoleophilia genomes. In contrast, 24 glycopeptide-resistance proteins belonging to the *vanABFGHMOTXW* repertoire showed substantial sequence similarity to proteins from 355 reference genomes, representing approximately 35% of the Thermoleophilia genomes examined. These homologs were distributed across multiple taxonomic orders, including UBA2241, RBG-16-64-13, and BMS3ABIN01.

**Figure 5.**
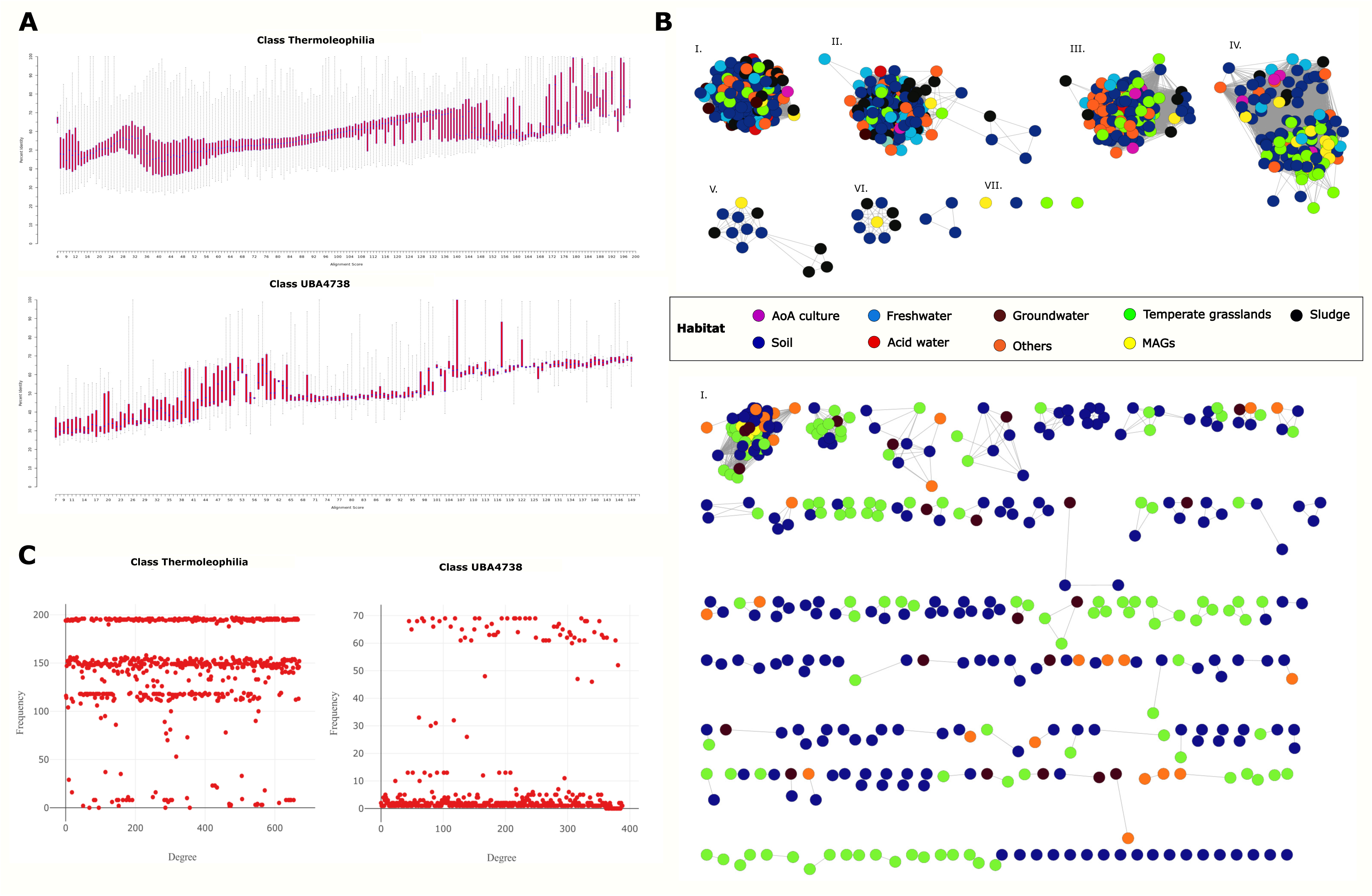
Sequence similarity networks (SSN) of ARGs in Thermoleophilia and UBA4738. (A) A boxplot showing distribution of alignment scores of 671 and 389 sequences of Thermoleophilia and UBA4738 classes against matching reference sequences. (B) The SSN of all ARGs (671 nodes) in Thermoleophilia from several habitats formed 7 different clusters (our MAGs are shown in yellow; n=24). For UBA4738, the SSN forms 1 cluster with MAGs encoding for *vanT,* whereas the other 389 nodes in UBA4738 form 126 connected components. (C) The degree distribution graph showing the frequency of a node occurring in a single cluster in both classes.

The Thermoleophilia ARG sequences formed a network containing 671 nodes and 11 connected components, including seven larger isofunctional clusters and four isolated nodes (Figures 5B and 5C). The query sequences were distributed among seven groups, with *vanT*, *vanOG*, and *vanH* forming clusters associated with vancomycin-resistance determinants, while *vanW* and *vanX* formed clusters associated with glycopeptide-resistance determinants including teicoplanin resistance. The *vanAFMB* cluster contained sequences associated with both vancomycin- and teicoplanin-resistance systems. These patterns indicate that several glycopeptide-resistance determinants identified in the Thermoleophilia MAGs are closely related to homologs occurring across diverse reference genomes. A similar analysis of four *vanT* sequences from UBA4738 revealed substantial sequence similarity to homologs from 67 reference genomes, representing approximately 74% of the UBA4738 genomes examined. The resulting SSN contained 389 nodes distributed among 127 connected components, including one larger isofunctional cluster and 17 isolated nodes (Figure 5C). All four UBA4738 query sequences mapped to the same major cluster, indicating strong sequence conservation among these *vanT* proteins and their reference homologs.

## Discussion

### Metabolic flexibility of Actinobacteria in hydrothermal steam vents

Actinobacteria are renowned for their metabolic diversity and capacity to produce structurally diverse secondary metabolites, including a large proportion of clinically important antibiotics as well as anticancer, antifungal, and other bioactive compounds (58). Despite this well-established biotechnological importance, the ecological roles and metabolic capabilities of Actinobacteria inhabiting geothermally active subsurface environments remain comparatively poorly characterized. Here, genome-resolved metagenomic analysis of Hawaiian hydrothermal steam-vent communities recovered 58 Actinobacterial MAGs representing five classes: Acidimicrobiia, Actinomycetia, CALGFH01, Thermoleophilia, and UBA4738. Comparative genomic analyses revealed broad metabolic versatility across these lineages, including pathways associated with central carbon metabolism, amino acid biosynthesis and degradation, lipid metabolism, carbon monoxide and sulfur metabolism, nutrient cycling, secondary metabolite biosynthesis, and antibiotic resistance. Collectively, these findings suggest that Actinobacteria may occupy diverse metabolic niches within the chemically heterogeneous environments of hydrothermal steam vents.

The metabolic profiles of the MAGs indicate that multiple lineages possess the potential to use both organic and inorganic substrates for energy and biomass production. Members of CALGFH01 were particularly notable for the presence of pathways associated with carbon fixation and inorganic-carbon metabolism, including the reductive tricarboxylic acid cycle. The occurrence of these pathways, together with genes involved in utilization of inorganic electron donors, is consistent with the potential for chemolithotrophic or mixotrophic growth in this lineage. However, the presence and completeness of metabolic pathways in MAGs provide evidence of physiological potential rather than direct evidence of activity *in situ*. Experimental or transcriptomic studies will therefore be required to establish the contribution of these pathways to energy conservation and carbon fixation in hydrothermal environments.

Several additional metabolic features may contribute to persistence under the physicochemically heterogeneous conditions of steam vents. The widespread occurrence of pathways for amino acid and nucleotide biosynthesis indicates substantial capacity for autonomous biomass production, while the abundance of acyl-CoA dehydrogenases suggests that fatty-acid degradation may provide an important source of reducing equivalents and acetyl-CoA. The detection of ectoine biosynthesis in the Pseudonocardia-affiliated MAG S34-36_bin.43 is also noteworthy because ectoine is an established compatible solute involved in protection against osmotic and other environmental stresses (59, 60). Such stress-response capabilities may be advantageous in steam-vent habitats characterized by steep gradients in temperature, moisture, salinity, and chemical composition.

The detection of carbon monoxide oxidation genes provides another potential mechanism for energy acquisition. The *coxSML* genes encode components of aerobic or microaerobic carbon monoxide dehydrogenase systems and have been associated with CO oxidation across diverse bacterial lineages (61–64). Carbon monoxide can serve as an energy and carbon source for carboxydotrophic microorganisms, particularly in environments where organic carbon is limited (65–67). The occurrence of *coxSML* across MAGs representing all five Actinobacterial classes therefore suggests that CO oxidation may represent a previously underappreciated metabolic capability in hydrothermal steam-vent Actinobacteria. Rather than indicating that CO is necessarily a dominant energy source, these findings suggest that CO oxidation could provide an additional metabolic option under carbon- or energy-limited conditions. Determining whether these organisms actively oxidize CO *in situ* will require physiological and gene expression-based analyses.

The distribution of sulfur-associated genes further supports the metabolic flexibility of these organisms. Genes such as *metE*, *ahcY*, and *cysH* indicate potential for transformation and assimilation of sulfur-containing compounds, while *dmoB* and other associated genes suggest the capacity to utilize reduced organosulfur compounds. These functions could provide both metabolic intermediates and reducing equivalents while linking organic sulfur metabolism to the broader sulfur cycle (68–75). Thus, the coexistence of heterotrophic pathways with pathways for utilization of inorganic or reduced carbon and sulfur compounds may allow Actinobacteria to exploit multiple energy sources as environmental conditions fluctuate.

### Potential contributions to biogeochemical cycling

The distribution of nutrient-cycling genes across the MAGs suggests that Actinobacteria may contribute to several elemental cycles within hydrothermal steam-vent communities. The MEBS analysis indicated substantial variation among taxonomic classes in the representation of carbon, nitrogen, sulfur, oxygen, and iron cycling functions. CALGFH01 exhibited relatively high entropy scores for nitrogen, oxygen, and iron-associated functions, whereas UBA4738 showed comparatively high sulfur-cycling scores. These differences are consistent with metabolic differentiation among lineages and may reflect adaptation to distinct chemical niches within the steam-vent environment.

Nitrogen-cycling potential was widespread across the dataset. The presence of *nirBD* in all five classes indicates a broad capacity for reduction of nitrite to ammonium, whereas nitrogen-fixation pathways were comparatively more complete in CALGFH01. Nitrite reduction can contribute to nitrogen assimilation and, under appropriate physiological conditions, to nitrogen conservation or dissimilatory nitrogen metabolism (76–82). The distribution of these pathways suggests that Actinobacteria may participate in nitrogen transformations where nitrogen availability and redox conditions fluctuate. Nevertheless, pathway completeness alone does not establish whether individual MAGs perform assimilatory or dissimilatory reactions in the environment, and these alternative functions should be distinguished experimentally.

Sulfur transformations were also prominent. The occurrence of sulfur oxidation and reduction-associated pathways, including *dsrEFHABCKMJOP*, *apt/sat*, *sor*, and other sulfur-associated gene systems, particularly in UBA4738 and CALGFH01, suggests that these lineages may interact with the diverse reduced and oxidized sulfur compounds generated within hydrothermal systems. Such metabolic capabilities could be advantageous across the steep redox gradients that characterize steam-vent habitats. Similarly, genes associated with iron reduction and Fe(II) oxidation were differentially represented among the classes, with Thermoleophilia showing greater representation of iron-reduction potential and CALGFH01 showing greater representation of Fe(II)-oxidation-associated functions. These lineage-specific patterns suggest that different Actinobacterial groups may contribute to distinct portions of the local redox network.

The broad distribution of nutrient-cycling functions is particularly relevant to hydrothermal steam vents because these systems generate strong chemical gradients over relatively small spatial scales. Reduced compounds originating from geothermal fluids can encounter oxidized compounds introduced from the atmosphere, surface waters, or surrounding soils, creating numerous potential electron donor–acceptor combinations. The metabolic diversity observed across the MAGs may therefore allow Actinobacteria to exploit multiple redox conditions and contribute to elemental transformations within steam-vent biofilms. Importantly, these interpretations remain based on genomic potential; measurements of gene expression, metabolite fluxes, and in situ geochemical gradients will be necessary to determine the actual contribution of these organisms to biogeochemical cycling.

### Secondary metabolite biosynthetic potential

The extensive diversity of biosynthetic gene clusters (BGCs) identified across the 58 MAGs highlights the substantial secondary-metabolite potential of Actinobacteria inhabiting hydrothermal steam vents. Genome mining identified 219 BGCs associated with terpenes, RiPPs, NRPSs, PKSs, β-lactones, and several less abundant biosynthetic classes. The predominance of terpene and RiPP-associated clusters, together with the substantial representation of NRPS and PKS pathways, indicates that secondary metabolism is a prominent feature of these genomes. This observation is consistent with the broader importance of Actinobacteria as prolific producers of structurally diverse natural products (83–87).

The distribution of BGC classes varied substantially among phylogenetic groups. Actinomycetia contained a large proportion of the NRPS-associated clusters, whereas CALGFH01 was enriched in PKS-associated BGCs. RiPP-associated clusters were particularly represented in Thermoleophilia, Acidimicrobiia, and Actinomycetia. Several less common BGC classes were also detected, including type III polyketide synthases and HglE-KS-associated clusters in CALGFH01, RRE-containing and type II polyketide synthase clusters in Thermoleophilia, and arylpolyene- and ranthipeptide-associated clusters in Acidimicrobiia. These lineage-specific patterns suggest that secondary metabolism may have diversified differently among the major Actinobacterial groups inhabiting these environments. The ecological functions of these biosynthetic pathways are potentially diverse. Secondary metabolites can mediate microbial competition, signaling, metal acquisition, oxidative-stress protection, and interactions with other organisms (88–96). For example, arylpolyene biosynthetic pathways have been associated with antioxidant functions, whereas peptide and polyketide products can contribute to microbial antagonism and environmental adaptation. The occurrence of these pathways in hydrothermal steam-vent Actinobacteria therefore raises the possibility that secondary metabolism contributes to survival and competition under extreme environmental conditions.

However, BGC identification does not establish expression or production of the predicted metabolites. Likewise, the presence of a BGC cannot by itself establish its ecological function. Consequently, the BGCs identified here should be considered a reservoir of biosynthetic potential rather than evidence of active secondary-metabolite production. Metatranscriptomic, metabolomic, and experimental studies will be required to determine which clusters are expressed and to characterize their products. Nevertheless, the diversity of BGCs recovered from these previously undercharacterized Actinobacterial lineages demonstrates that hydrothermal steam vents represent a promising source of unexplored microbial biosynthetic diversity.

### Glycopeptide-resistance determinants in hydrothermal Actinobacteria

Antibiotic resistance genes were detected in 50 of the 58 MAGs, indicating that resistance-associated functions are widespread among the Actinobacteria recovered from Hawaiian hydrothermal steam vents. The predominance of glycopeptide-resistance-associated determinants was particularly notable, with *van*-associated genes occurring across multiple phylogenetic classes and Thermoleophilia exhibiting the highest overall ARG representation. This observation expands the known ecological distribution of glycopeptide-resistance-associated genes beyond the clinical and antibiotic-producing microorganisms in which these systems have been most extensively studied.

Glycopeptide antibiotics such as vancomycin and teicoplanin inhibit peptidoglycan synthesis by binding to the D-Ala-D-Ala terminus of cell-wall precursors. Established resistance systems alter this target, most commonly through replacement of D-Ala-D-Ala with D-Ala-D-Lac or D-Ala-D-Ser, thereby substantially reducing glycopeptide binding (97–102). The occurrence of multiple components of these systems – including *vanA/B*, *vanH*, *vanX*, *vanT*, and *vanW* – in the Actinobacterial MAGs suggests that some of these organisms possess substantial genomic potential for glycopeptide resistance. In particular, the concentration of *van*-associated genes in Thermoleophilia is notable because this class is relatively poorly characterized compared with medically important Actinobacterial groups. It is also intriguing to note that vancomycin resistance related genes were detected previously in the Hawaii steam vent-associated early-branching cyanobacterium *Gloeobacter kilaueensis* JS1, which was isolated in 2013 (103).

The ecological origin and function of these resistance determinants remain unresolved. Hydrothermal steam vents are not expected to experience the same direct antibiotic selection pressures as clinical environments, raising the possibility that some resistance-associated genes represent intrinsic or ancestral cellular functions rather than adaptations to contemporary antibiotic exposure. Alternatively, resistance determinants may have been acquired through horizontal gene transfer from other environmental or antibiotic-producing microorganisms. Antibiotic-producing Actinobacteria are themselves known to carry resistance mechanisms that protect against their own bioactive compounds, providing a potential evolutionary reservoir of resistance determinants (104, 105). Distinguishing among these possibilities will require gene-specific phylogenetic analyses, examination of genomic context and gene-cluster architecture, and comparative analysis with experimentally characterized resistance loci.

The widespread detection of *van*-associated genes should also be interpreted cautiously. Identification of ARGs in MAGs demonstrates genomic potential for resistance but does not establish that the corresponding organisms are phenotypically resistant to vancomycin or teicoplanin. In particular, the functional contribution of individual genes such as *vanW* remains incompletely resolved (106). Demonstrating resistance would require expression and physiological assays, ideally including susceptibility testing and characterization of the relevant peptidoglycan precursors. Nevertheless, the distribution of multiple components of glycopeptide-resistance systems across these environmental MAGs warrants further investigation.

### Sequence similarity reveals conservation of glycopeptide-resistance-associated proteins

Sequence similarity network analysis provided additional evidence that several glycopeptide-resistance-associated proteins identified in Thermoleophilia and UBA4738 are related to homologous proteins occurring across diverse reference genomes. In Thermoleophilia, *vanT*, *vanH*, *vanX*, *vanW*, and *vanAFMB*-associated sequences formed distinct sequence-similarity groups, whereas the *rphA/rphB* sequences showed more limited similarity to reference proteins. The Thermoleophilia query sequences were distributed among multiple network components and clustered with homologs from other taxonomic groups, indicating that these resistance-associated proteins are members of broader protein families rather than highly divergent sequences restricted to the hydrothermal MAGs. The UBA4738 analysis similarly showed that the four MAG-derived *vanT* sequences clustered with homologous proteins from a substantial fraction of the available UBA4738 reference genomes. The shared sequence similarity among these proteins is consistent with conservation of *vanT*-associated functions across members of this lineage. Together, the Thermoleophilia and UBA4738 networks suggest that glycopeptide-resistance-associated protein families are more broadly distributed among environmental Actinobacteria than previously recognized.

The sequence networks, however, do not by themselves resolve the evolutionary mechanisms responsible for this conservation. Sequence similarity cannot distinguish vertical inheritance from horizontal gene transfer or differential gene loss, and network topology alone cannot establish positive or negative selection. Likewise, sequence similarity does not demonstrate that differences among proteins alter antibiotic susceptibility. Phylogenetic reconstruction of individual *van* genes, analysis of their genomic neighborhoods, and comparison of gene-cluster organization will be necessary to determine the evolutionary history and selective forces shaping these resistance determinants. Despite these limitations, the occurrence of conserved glycopeptide-resistance-associated proteins in previously undercharacterized environmental Actinobacteria provides an important starting point for investigating the evolution and ecological function of antibiotic resistance outside clinical environments. Structural and biochemical characterization of divergent or lineage-specific proteins may ultimately reveal functional differences that cannot be resolved through sequence similarity alone.

## Conclusions

This study expands the genomic characterization of Actinobacteria inhabiting geothermally active Hawaiian steam vents and reveals substantial functional diversity across five phylogenetic classes. The 58 MAGs collectively encode metabolic pathways capable of supporting heterotrophic growth, utilization of complex organic substrates, and potential chemolithotrophic energy acquisition through carbon monoxide and sulfur metabolism. Their diverse nitrogen-, sulfur-, and iron-associated functions further suggest that these organisms may contribute to biogeochemical transformations within chemically heterogeneous steam-vent communities. In parallel, the extensive diversity of secondary-metabolite BGCs highlights the biosynthetic potential of these environments, while the widespread occurrence of ARGs (particularly glycopeptide-resistance-associated determinants in Thermoleophilia) reveals an additional dimension of Actinobacterial functional diversity. Sequence similarity networks indicate that several of these resistance-associated proteins are conserved within broader environmental protein families, although their evolutionary origins and physiological roles remain to be resolved. Overall, these findings suggest that hydrothermal steam vents harbor previously underrecognized Actinobacterial diversity with the genomic potential to participate in energy acquisition, elemental cycling, microbial interactions, secondary metabolism, and antibiotic resistance. Future integration of genome-resolved metagenomics with metatranscriptomics, metabolomics, geochemical measurements, and experimental characterization will be essential for determining which of these predicted functions are active in situ and for establishing how Actinobacteria contribute to the ecology and evolution of geothermally active terrestrial ecosystems.

## Acknowledgements

We gratefully acknowledge the computational resources provided on the high-performance computing cluster operated by Research Technology Services at The George Washington University.

## Data Availability

All raw data used in this study are publicly available at European Nucleotide Archives (ENA) under the project accession number PRJEB52128. MAGs used in this study have been deposited to Zenodo at this DOI number: 10.5281/zenodo.22727583.

## Author Contribution (CRediT)

**Shekhar Nagar**: investigation, formal analysis, data visualization, writing – original draft preparation. **Chengxuan Zhang**: formal analysis, writing – review and editing. **Jimmy H. Saw**: conceptualization, supervision, resources, funding acquisition, data curation, project administration, writing – review and editing.

## References

1. Gomez-Alvarez V, King GM, Nüsslein K. 2007. Comparative bacterial diversity in recent Hawaiian volcanic deposits of different ages: Microbial diversity of four Hawaiian volcanic deposits. FEMS Microbiol Ecol 60:60–73.

2. Northup DE, Lavoie KH. 2015. 8. Microbial Diversity and Ecology of Lava Caves, p. 161–192. In Summers Engel, A (ed.), Microbial Life of Cave Systems. DE GRUYTER.

3. Costello EK, Halloy SRP, Reed SC, Sowell P, Schmidt SK. 2009. Fumarole-Supported Islands of Biodiversity within a Hyperarid, High-Elevation Landscape on Socompa Volcano, Puna de Atacama, Andes. Appl Environ Microbiol 75:735–747.

4. Wall K, Cornell J, Bizzoco RW, Kelley ST. 2015. Biodiversity hot spot on a hot spot: novel extremophile diversity in Hawaiian fumaroles. MicrobiologyOpen 4:267–281.

5. Riquelme C, Rigal F, Hathaway JJM, Northup DE, Spilde MN, Borges PAV, Gabriel R, Amorim IR, Dapkevicius MDLNE. 2015. Cave microbial community composition in oceanic islands: disentangling the effect of different colored mats in diversity patterns of Azorean lava caves. FEMS Microbiol Ecol 91:fiv141.

6. Barton HA. 2015. 4. Starving Artists: Bacterial Oligotrophic Heterotrophy in Caves, p. 79–104. In Summers Engel, A (ed.), Microbial Life of Cave Systems. DE GRUYTER.

7. Miller AZ, Pereira MFC, Calaforra JM, Forti P, Dionísio A, Saiz-Jimenez C. 2014. Siliceous Speleothems and Associated Microbe-Mineral Interactions from Ana Heva Lava Tube in Easter Island (Chile). Geomicrobiol J 31:236–245.

8. Prescott RD, Zamkovaya T, Donachie SP, Northup DE, Medley JJ, Monsalve N, Saw JH, Decho AW, Chain PSG, Boston PJ. 2022. Islands Within Islands: Bacterial Phylogenetic Structure and Consortia in Hawaiian Lava Caves and Fumaroles. Front Microbiol 13:934708.

9. Cockell CS, Santomartino R, Finster K, Waajen AC, Eades LJ, Moeller R, Rettberg P, Fuchs FM, Van Houdt R, Leys N, Coninx I, Hatton J, Parmitano L, Krause J, Koehler A, Caplin N, Zuijderduijn L, Mariani A, Pellari SS, Carubia F, Luciani G, Balsamo M, Zolesi V, Nicholson N, Loudon C-M, Doswald-Winkler J, Herová M, Rattenbacher B, Wadsworth J, Craig Everroad R, Demets R. 2020. Space station biomining experiment demonstrates rare earth element extraction in microgravity and Mars gravity. Nat Commun 11:5523.

10. Miao V, Davies J. 2010. Actinobacteria: the good, the bad, and the ugly. Antonie Van Leeuwenhoek 98:143–150.

11. Zaremba-Niedźwiedzka K, Andersson SGE. 2013. No Ancient DNA Damage in Actinobacteria from the Neanderthal Bone. PLoS ONE 8:e62799.

12. Dentzien-Dias P, Poinar G, Francischini H. 2017. A new actinomycete from a Guadalupian vertebrate coprolite from Brazil. Hist Biol 29:770–776.

13. Barton HA, Giarrizzo JG, Suarez P, Robertson CE, Broering MJ, Banks ED, Vaishampayan PA, Venkateswaran K. 2014. Microbial diversity in a Venezuelan orthoquartzite cave is dominated by the Chloroflexi (Class Ktedonobacterales) and Thaumarchaeota Group I.1c. Front Microbiol 5.

14. Hathaway JJM, Garcia MG, Balasch MM, Spilde MN, Stone FD, Dapkevicius MDLNE, Amorim IR, Gabriel R, Borges PAV, Northup DE. 2014. Comparison of Bacterial Diversity in Azorean and Hawai’ian Lava Cave Microbial Mats. Geomicrobiol J 31:205–220.

15. Jiang X, Ellabaan MMH, Charusanti P, Munck C, Blin K, Tong Y, Weber T, Sommer MOA, Lee SY. 2017. Dissemination of antibiotic resistance genes from antibiotic producers to pathogens. Nat Commun 8:15784.

16. Lund D, Coertze RD, Parras-Moltó M, Berglund F, Flach C-F, Johnning A, Larsson DGJ, Kristiansson E. 2023. Extensive screening reveals previously undiscovered aminoglycoside resistance genes in human pathogens. Commun Biol 6:812.

17. Jose PA, Jebakumar SRD. 2013. Non-streptomycete actinomycetes nourish the current microbial antibiotic drug discovery. Front Microbiol 4:240.

18. Hill P, Krištůfek V, Dijkhuizen L, Boddy C, Kroetsch D, Van Elsas JD. 2011. Land Use Intensity Controls Actinobacterial Community Structure. Microb Ecol 61:286–302.

19. Beattie AJ, Hay M, Magnusson B, De Nys R, Smeathers J, Vincent JFV. 2011. Ecology and bioprospecting. Austral Ecol 36:341–356.

20. Saw JH, Shlafstein MD, Pavloudi C, Monsalve N, Prescott RD, Chain PSG, Decho AW, Donachie SP. 2026. Amplicon and metagenomic data from fumarole-associated geothermal features of Hawaiʻi. Sci Data 10.1038/s41597-026-07734-x.

21. Nurk S, Meleshko D, Korobeynikov A, Pevzner PA. 2017. metaSPAdes: a new versatile metagenomic assembler. Genome Res 27:824–834.

22. Smith HE, Yun S. 2017. Evaluating alignment and variant-calling software for mutation identification in C. elegans by whole-genome sequencing. PLOS ONE 12:e0174446.

23. Kang DD, Li F, Kirton E, Thomas A, Egan R, An H, Wang Z. 2019. MetaBAT 2: an adaptive binning algorithm for robust and efficient genome reconstruction from metagenome assemblies. PeerJ 7:e7359.

24. The Genome Standards Consortium, Bowers RM, Kyrpides NC, Stepanauskas R, Harmon-Smith M, Doud D, Reddy TBK, Schulz F, Jarett J, Rivers AR, Eloe-Fadrosh EA, Tringe SG, Ivanova NN, Copeland A, Clum A, Becraft ED, Malmstrom RR, Birren B, Podar M, Bork P, Weinstock GM, Garrity GM, Dodsworth JA, Yooseph S, Sutton G, Glöckner FO, Gilbert JA, Nelson WC, Hallam SJ, Jungbluth SP, Ettema TJG, Tighe S, Konstantinidis KT, Liu W-T, Baker BJ, Rattei T, Eisen JA, Hedlund B, McMahon KD, Fierer N, Knight R, Finn R, Cochrane G, Karsch-Mizrachi I, Tyson GW, Rinke C, Lapidus A, Meyer F, Yilmaz P, Parks DH, Murat Eren A, Schriml L, Banfield JF, Hugenholtz P, Woyke T. 2017. Minimum information about a single amplified genome (MISAG) and a metagenome-assembled genome (MIMAG) of bacteria and archaea. Nat Biotechnol 35:725–731.

25. Bowers RM, Kyrpides NC, Stepanauskas R, Harmon-Smith M, Doud D, Reddy TBK, Schulz F, Jarett J, Rivers AR, Eloe-Fadrosh EA, Tringe SG, Ivanova NN, Copeland A, Clum A, Becraft ED, Malmstrom RR, Birren B, Podar M, Bork P, Weinstock GM, Garrity GM, Dodsworth JA, Yooseph S, Sutton G, Glöckner FO, Gilbert JA, Nelson WC, Hallam SJ, Jungbluth SP, Ettema TJG, Tighe S, Konstantinidis KT, Liu W-T, Baker BJ, Rattei T, Eisen JA, Hedlund B, McMahon KD, Fierer N, Knight R, Finn R, Cochrane G, Karsch-Mizrachi I, Tyson GW, Rinke C, Genome Standards Consortium, Lapidus A, Meyer F, Yilmaz P, Parks DH, Eren AM, Schriml L, Banfield JF, Hugenholtz P, Woyke T. 2017. Minimum information about a single amplified genome (MISAG) and a metagenome-assembled genome (MIMAG) of bacteria and archaea. Nat Biotechnol 35:725–731.

26. Jain C, Rodriguez-R LM, Phillippy AM, Konstantinidis KT, Aluru S. 2018. High throughput ANI analysis of 90K prokaryotic genomes reveals clear species boundaries. Nat Commun 9:5114.

27. Chaumeil P-A, Mussig AJ, Hugenholtz P, Parks DH. 2022. GTDB-Tk v2: memory friendly classification with the genome taxonomy database. Bioinformatics 38:5315–5316.

28. Chaumeil P-A, Mussig AJ, Hugenholtz P, Parks DH. 2019. GTDB-Tk: a toolkit to classify genomes with the Genome Taxonomy Database. Bioinforma Oxf Engl 10.1093/bioinformatics/btz848.

29. Parks DH, Chuvochina M, Rinke C, Mussig AJ, Chaumeil P-A, Hugenholtz P. 2022. GTDB: an ongoing census of bacterial and archaeal diversity through a phylogenetically consistent, rank normalized and complete genome-based taxonomy. Nucleic Acids Res 50:D785–D794.

30. Buchfink B, Reuter K, Drost H-G. 2021. Sensitive protein alignments at tree-of-life scale using DIAMOND. Nat Methods 18:366–368.

31. Edgar RC. 2004. MUSCLE: a multiple sequence alignment method with reduced time and space complexity. BMC Bioinformatics 5:113.

32. Price MN, Dehal PS, Arkin AP. 2010. FastTree 2 – Approximately Maximum-Likelihood Trees for Large Alignments. PLoS ONE 5:e9490.

33. Seemann T. 2014. Prokka: rapid prokaryotic genome annotation. Bioinformatics 30:2068–2069.

34. Pundir S, Martin MJ, O’Donovan C, The UniProt Consortium. 2016. UniProt Tools. Curr Protoc Bioinforma 53.

35. Asnicar F, Thomas AM, Beghini F, Mengoni C, Manara S, Manghi P, Zhu Q, Bolzan M, Cumbo F, May U, Sanders JG, Zolfo M, Kopylova E, Pasolli E, Knight R, Mirarab S, Huttenhower C, Segata N. 2020. Precise phylogenetic analysis of microbial isolates and genomes from metagenomes using PhyloPhlAn 3.0. Nat Commun 11:2500.

36. Letunic I, Bork P. 2021. Interactive Tree Of Life (iTOL) v5: an online tool for phylogenetic tree display and annotation. Nucleic Acids Res 49:W293–W296.

37. Hyatt D, Chen G-L, LoCascio PF, Land ML, Larimer FW, Hauser LJ. 2010. Prodigal: prokaryotic gene recognition and translation initiation site identification. BMC Bioinformatics 11:119.

38. Zhou Z, Tran PQ, Breister AM, Liu Y, Kieft K, Cowley ES, Karaoz U, Anantharaman K. 2022. METABOLIC: high-throughput profiling of microbial genomes for functional traits, metabolism, biogeochemistry, and community-scale functional networks. Microbiome 10:33.

39. Kanehisa M, Sato Y. 2020. KEGG Mapper for inferring cellular functions from protein sequences. Protein Sci 29:28–35.

40. Aramaki T, Blanc-Mathieu R, Endo H, Ohkubo K, Kanehisa M, Goto S, Ogata H. 2020. KofamKOALA: KEGG Ortholog assignment based on profile HMM and adaptive score threshold. Bioinformatics 36:2251–2252.

41. Finn RD, Bateman A, Clements J, Coggill P, Eberhardt RY, Eddy SR, Heger A, Hetherington K, Holm L, Mistry J, Sonnhammer ELL, Tate J, Punta M. 2014. Pfam: the protein families database. Nucleic Acids Res 42:D222–D230.

42. Haft DH. 2003. The TIGRFAMs database of protein families. Nucleic Acids Res 31:371–373.

43. R Core Team. 2021. R Statistical Software.

44. Zhang H, Yohe T, Huang L, Entwistle S, Wu P, Yang Z, Busk PK, Xu Y, Yin Y. 2018. dbCAN2: a meta server for automated carbohydrate-active enzyme annotation. Nucleic Acids Res 46:W95–W101.

45. Rawlings ND, Morton FR, Kok CY, Kong J, Barrett AJ. 2007. MEROPS: the peptidase database. Nucleic Acids Res 36:D320–D325.

46. De Anda V, Zapata-Peñasco I, Poot-Hernandez AC, Eguiarte LE, Contreras-Moreira B, Souza V. 2017. MEBS, a software platform to evaluate large (meta)genomic collections according to their metabolic machinery: unraveling the sulfur cycle. GigaScience 6.

47. Blin K, Shaw S, Augustijn HE, Reitz ZL, Biermann F, Alanjary M, Fetter A, Terlouw BR, Metcalf WW, Helfrich EJN, van Wezel GP, Medema MH, Weber T. 2023. antiSMASH 7.0: new and improved predictions for detection, regulation, chemical structures and visualisation. Nucleic Acids Res 51:W46–W50.

48. McArthur AG, Waglechner N, Nizam F, Yan A, Azad MA, Baylay AJ, Bhullar K, Canova MJ, De Pascale G, Ejim L, Kalan L, King AM, Koteva K, Morar M, Mulvey MR, O’Brien JS, Pawlowski AC, Piddock LJV, Spanogiannopoulos P, Sutherland AD, Tang I, Taylor PL, Thaker M, Wang W, Yan M, Yu T, Wright GD. 2013. The Comprehensive Antibiotic Resistance Database. Antimicrob Agents Chemother 57:3348–3357.

49. Alcock BP, Huynh W, Chalil R, Smith KW, Raphenya AR, Wlodarski MA, Edalatmand A, Petkau A, Syed SA, Tsang KK, Baker SJC, Dave M, McCarthy MC, Mukiri KM, Nasir JA, Golbon B, Imtiaz H, Jiang X, Kaur K, Kwong M, Liang ZC, Niu KC, Shan P, Yang JYJ, Gray KL, Hoad GR, Jia B, Bhando T, Carfrae LA, Farha MA, French S, Gordzevich R, Rachwalski K, Tu MM, Bordeleau E, Dooley D, Griffiths E, Zubyk HL, Brown ED, Maguire F, Beiko RG, Hsiao WWL, Brinkman FSL, Van Domselaar G, McArthur AG. 2023. CARD 2023: expanded curation, support for machine learning, and resistome prediction at the Comprehensive Antibiotic Resistance Database. Nucleic Acids Res 51:D690–D699.

50. Bortolaia V, Kaas RS, Ruppe E, Roberts MC, Schwarz S, Cattoir V, Philippon A, Allesoe RL, Rebelo AR, Florensa AF, Fagelhauer L, Chakraborty T, Neumann B, Werner G, Bender JK, Stingl K, Nguyen M, Coppens J, Xavier BB, Malhotra-Kumar S, Westh H, Pinholt M, Anjum MF, Duggett NA, Kempf I, Nykäsenoja S, Olkkola S, Wieczorek K, Amaro A, Clemente L, Mossong J, Losch S, Ragimbeau C, Lund O, Aarestrup FM. 2020. ResFinder 4.0 for predictions of phenotypes from genotypes. J Antimicrob Chemother 75:3491–3500.

51. Yin X, Jiang X-T, Chai B, Li L, Yang Y, Cole JR, Tiedje JM, Zhang T. 2018. ARGs-OAP v2.0 with an expanded SARG database and Hidden Markov Models for enhancement characterization and quantification of antibiotic resistance genes in environmental metagenomes. Bioinformatics 34:2263–2270.

52. Krzywinski M, Schein J, Birol İ, Connors J, Gascoyne R, Horsman D, Jones SJ, Marra MA. 2009. Circos: An information aesthetic for comparative genomics. Genome Res 19:1639–1645.

53. Fu L, Niu B, Zhu Z, Wu S, Li W. 2012. CD-HIT: accelerated for clustering the next-generation sequencing data. Bioinformatics 28:3150–3152.

54. Gerlt JA, Bouvier JT, Davidson DB, Imker HJ, Sadkhin B, Slater DR, Whalen KL. 2015. Enzyme Function Initiative-Enzyme Similarity Tool (EFI-EST): A web tool for generating protein sequence similarity networks. Biochim Biophys Acta BBA - Proteins Proteomics 1854:1019–1037.

55. Otasek D, Morris JH, Bouças J, Pico AR, Demchak B. 2019. Cytoscape Automation: empowering workflow-based network analysis. Genome Biol 20:185.

56. O’Leary NA, Cox E, Holmes JB, Anderson WR, Falk R, Hem V, Tsuchiya MTN, Schuler GD, Zhang X, Torcivia J, Ketter A, Breen L, Cothran J, Bajwa H, Tinne J, Meric PA, Hlavina W, Schneider VA. 2024. Exploring and retrieving sequence and metadata for species across the tree of life with NCBI Datasets. Sci Data 11:732.

57. Yao R, Schneider E, Ryan TJ, Galivan J. 1996. Human gamma-glutamyl hydrolase: cloning and characterization of the enzyme expressed in vitro. Proc Natl Acad Sci 93:10134–10138.

58. Hopwood DA. 2007. Streptomyces in nature and medicine: the antibiotic makers. Oxford University Press.

59. Abdel-Aziz H, Wadie W, Abdallah DM, Lentzen G, Khayyal MT. 2013. Novel effects of ectoine, a bacteria-derived natural tetrahydropyrimidine, in experimental colitis. Phytomedicine 20:585–591.

60. Camarasa C, Faucet V, Dequin S. 2007. Role in anaerobiosis of the isoenzymes for *Saccharomyces cerevisiae* fumarate reductase encoded by *OSM1* and *FRDS1*. Yeast 24:391– 401.

61. Meyer O, Schlegel HG. 1983. BIOLOGY OF AEROBIC CARBON MONOXIDE-OXIDIZING BACTERIA. Annu Rev Microbiol 37:277–310.

62. Gadkari D, Schricker K, Acker G, Kroppenstedt RM, Meyer O. 1990. Streptomyces thermoautotrophicus sp. nov., a Thermophilic CO- and H2-Oxidizing Obligate Chemolithoautotroph. Appl Environ Microbiol 56:3727–3734.

63. Dunfield KE, King GM. 2004. Molecular Analysis of Carbon Monoxide-Oxidizing Bacteria Associated with Recent Hawaiian Volcanic Deposits. Appl Environ Microbiol 70:4242–4248.

64. Hille R. 2005. Molybdenum-containing hydroxylases. Arch Biochem Biophys 433:107–116.

65. Amend JP, McCollom TM, Hentscher M, Bach W. 2011. Catabolic and anabolic energy for chemolithoautotrophs in deep-sea hydrothermal systems hosted in different rock types. Geochim Cosmochim Acta 75:5736–5748.

66. King GM. 2003. Molecular andCulture-Based Analyses of Aerobic Carbon Monoxide OxidizerDiversity†. Appl Environ Microbiol 69:7257–7265.

67. Patrauchan MA, Miyazawa D, LeBlanc JC, Aiga C, Florizone C, Dosanjh M, Davies J, Eltis LD, Mohn WW. 2012. Proteomic Analysis of Survival of Rhodococcus jostii RHA1 during Carbon Starvation. Appl Environ Microbiol 78:6714–6725.

68. Luo W, Zhao M, Dwidar M, Gao Y, Xiang L, Wu X, Medema MH, Xu S, Li X, Schäfer H, Chen M, Feng R, Zhu Y. 2024. Microbial assimilatory sulfate reduction-mediated H2S: an overlooked role in Crohn’s disease development. Microbiome 12:152.

69. Nagar S, Talwar C, Motelica-Heino M, Richnow H-H, Shakarad M, Lal R, Negi RK. 2022. Microbial Ecology of Sulfur Biogeochemical Cycling at a Mesothermal Hot Spring Atop Northern Himalayas, India. Front Microbiol 13:848010.

70. Chiku T, Padovani D, Zhu W, Singh S, Vitvitsky V, Banerjee R. 2009. H2S Biogenesis by Human Cystathionine γ-Lyase Leads to the Novel Sulfur Metabolites Lanthionine and Homolanthionine and Is Responsive to the Grade of Hyperhomocysteinemia. J Biol Chem 284:11601–11612.

71. Kanagawa T, Kelly DP. 1986. Breakdown of dimethyl sulphide by mixed cultures and by Thiobacillus thioparus. FEMS Microbiol Lett 34:13–19.

72. Pol A, Op Den Camp HJM, Mees SGM, Kersten MASH, Van Der Drift C. 1994. Isolation of a dimethylsulfide-utilizingHyphomicrobium species and its application in biofiltration of polluted air. Biodegradation 5:105–112.

73. De Zwart J, Sluis J, Kuenen JG. 1997. Competition for Dimethyl Sulfide and Hydrogen Sulfide by Methylophaga sulfidovorans and Thiobacillus thioparus T5 in Continuous Cultures. Appl Environ Microbiol 63:3318–3322.

74. Boden R, Kelly DP, Murrell JC, Schäfer H. 2010. Oxidation of dimethylsulfide to tetrathionate by *Methylophaga thiooxidans* sp. nov.: a new link in the sulfur cycle. Environ Microbiol 12:2688–2699.

75. Koch T, Dahl C. 2018. A novel bacterial sulfur oxidation pathway provides a new link between the cycles of organic and inorganic sulfur compounds. ISME J 12:2479–2491.

76. Richardson DJ. 2000. Bacterial respiration: a flexible process for a changing environment 1999 Fleming Lecture (Delivered at the 144th meeting of the Society for General Microbiology, 8 September 1999). Microbiology 146:551–571.

77. Jørgensen H, Brandt K, Lauridsen C. 2008. Year rather than farming system influences protein utilization and energy value of vegetables when measured in a rat model. Nutr Res 28:866– 878.

78. Bakken LR, Bergaust L, Liu B, Frostegård Å. 2012. Regulation of denitrification at the cellular level: a clue to the understanding of N 2 O emissions from soils. Philos Trans R Soc B Biol Sci 367:1226–1234.

79. Maier RM. 2015. Biogeochemical Cycling, p. 339–373. In Environmental Microbiology. Elsevier.

80. Harborne NR, Griffiths L, Busby SJW, Cole JA. 1992. Transcriptional control, translation and function of the products of the five open reading frames of the *Escherichia coli nir* operon. Mol Microbiol 6:2805–2813.

81. Stewart V. 1994. Regulation of nitrate and nitrite reductase synthesis in enterobacteria. Antonie Van Leeuwenhoek 66:37–45.

82. Cole JA, Brown CM. 1980. NITRITE REDUCTION TO AMMONIA BY FERMENTATIVE BACTERIA: A SHORT CIRCUIT IN THE BIOLOGICAL NITROGEN CYCLE. FEMS Microbiol Lett 7:65–72.

83. Bérdy J. 2005. Bioactive Microbial Metabolites: A Personal View. J Antibiot (Tokyo) 58:1–26.

84. Vrancken K, Anné J. 2009. Secretory Production of Recombinant Proteins by *Streptomyces*. Future Microbiol 4:181–188.

85. Barka EA, Vatsa P, Sanchez L, Gaveau-Vaillant N, Jacquard C, Klenk H-P, Clément C, Ouhdouch Y, Van Wezel GP. 2016. Taxonomy, Physiology, and Natural Products of Actinobacteria. Microbiol Mol Biol Rev 80:1–43.

86. Bosi E, Taviani E, Avesani A, Doni L, Auguste M, Oliveri C, Leonessi M, Martinez-Urtaza J, Vetriani C, Vezzulli L. 2024. Pan-Genome Provides Insights into *Vibrio* Evolution and Adaptation to Deep-Sea Hydrothermal Vents. Genome Biol Evol 16:evae131.

87. Boddy CN. 2014. Bioinformatics tools for genome mining of polyketide and non-ribosomal peptides. J Ind Microbiol Biotechnol 41:443–450.

88. Ridley CP, Lee HY, Khosla C. 2008. Evolution of polyketide synthases in bacteria. Proc Natl Acad Sci 105:4595–4600.

89. Van Bergeijk DA, Terlouw BR, Medema MH, Van Wezel GP. 2020. Ecology and genomics of Actinobacteria: new concepts for natural product discovery. Nat Rev Microbiol 18:546–558.

90. Parra J, Beaton A, Seipke RF, Wilkinson B, Hutchings MI, Duncan KR. 2023. Antibiotics from rare actinomycetes, beyond the genus Streptomyces. Curr Opin Microbiol 76:102385.

91. Mashakhetri KD, Aishwarya CS, Prusty T, Bast F. 2024. Secondary Metabolites from Extremophiles, p. 177–201. In Shah, MP, Dey, S (eds.), Trends in Biotechnology of Polyextremophiles. Springer Nature Switzerland, Cham.

92. Nagar S, Bharti M, Negi RK. 2023. Genome-resolved metagenomics revealed metal-resistance, geochemical cycles in a Himalayan hot spring. Appl Microbiol Biotechnol 107:3273–3289.

93. Winkel-Shirley B. 2001. Flavonoid Biosynthesis. A Colorful Model for Genetics, Biochemistry, Cell Biology, and Biotechnology. Plant Physiol 126:485–493.

94. Austin MB, Noel JP. 2003. The chalcone synthase superfamily of type III polyketide synthases. Nat Prod Rep 20:79–110.

95. Ajesh BR, Sariga R, Nakkeeran S, Renukadevi P, Saranya N, Alkahtani S. 2024. Insights on mining the pangenome of Sphingobacterium thalpophilum NMS02 S296 from the resistant banana cultivar Pisang lilin confirms the antifungal action against Fusarium oxysporum f. sp. cubense. Front Microbiol 15:1443195.

96. Russell AH, Truman AW. 2020. Genome mining strategies for ribosomally synthesised and post-translationally modified peptides. Comput Struct Biotechnol J 18:1838–1851.

97. Bugg TDH, Wright GD, Dutka-Malen S, Arthur M, Courvalin P, Walsh CT. 1991. Molecular basis for vancomycin resistance in Enterococcus faecium BM4147: biosynthesis of a depsipeptide peptidoglycan precursor by vancomycin resistance proteins VanH and VanA. Biochemistry 30:10408–10415.

98. Arthur M, Molinas C, Bugg TD, Wright GD, Walsh CT, Courvalin P. 1992. Evidence for in vivo incorporation of D-lactate into peptidoglycan precursors of vancomycin-resistant enterococci. Antimicrob Agents Chemother 36:867–869.

99. Yushchuk O, Binda E, Marinelli F. 2020. Glycopeptide Antibiotic Resistance Genes: Distribution and Function in the Producer Actinomycetes. Front Microbiol 11:1173.

100. Reynolds PE, Depardieu F, Dutka-Malen S, Arthur M, Courvalin P. 1994. Glycopeptide resistance mediated by enterococcal transposon Tn 1546 requires production of VanX for hydrolysis of D-alanyl-D-alanine. Mol Microbiol 13:1065–1070.

101. Wu Z, Wright GD, Walsh CT. 1995. Overexpression, Purification, and Characterization of VanX, a D-, D-Dipeptidase which Is Essential for Vancomycin Resistance in Enterococcus faecium BM4147. Biochemistry 34:2455–2463.

102. Arthur M, Depardieu F, Cabanié L, Reynolds P, Courvalin P. 1998. Requirement of the VanY and VanX D, D-peptidases for glycopeptide resistance in enterococci. Mol Microbiol 30:819– 830.

103. Saw JHW, Schatz M, Brown MV, Kunkel DD, Foster JS, Shick H, Christensen S, Hou S, Wan X, Donachie SP. 2013. Cultivation and complete genome sequencing of Gloeobacter kilaueensis sp. nov., from a lava cave in Kīlauea Caldera, Hawai’i. PloS One 8:e76376.

104. Schäberle TF, Vollmer W, Frasch H-J, Hüttel S, Kulik A, Röttgen M, Von Thaler A-K, Wohlleben W, Stegmann E. 2011. Self-Resistance and Cell Wall Composition in the Glycopeptide Producer Amycolatopsis balhimycina. Antimicrob Agents Chemother 55:4283– 4289.

105. Marcone GL, Binda E, Carrano L, Bibb M, Marinelli F. 2014. Relationship between Glycopeptide Production and Resistance in the Actinomycete Nonomuraea sp. ATCC 39727. Antimicrob Agents Chemother 58:5191–5201.

106. Lebreton F, Cattoir V. 2019. Resistance to Glycopeptide Antibiotics, p. 51–80. In Bonev, BB, Brown, NM (eds.), Bacterial Resistance to Antibiotics – From Molecules to Man, 1st ed. Wiley.

